## Supplementary Information for "Integrated experimental-computational analysis of a liver-islet microphysiological system for human-centric diabetes research"

1 Drug Metabolism and Pharmacokinetics, Research and Early Development, Cardiovascular, Renal and Metabolism (CVRM), BioPharmaceuticals R&D, AstraZeneca, Gothenburg, Sweden.

2 Department of Biomedical Engineering, Linköping University, Linköping, Sweden.

3 TissUse GmbH, Berlin, Germany.

4 Bioscience, Research and Early Development, Cardiovascular, Renal and Metabolism (CVRM), BioPharmaceuticals R&D, AstraZeneca, Gothenburg, Sweden.

5 Center for Medical Image Science and Visualization (CMIV), Linköping University, Linköping, Sweden.

[†]Shared senior author

*Corresponding author

Gunnar Cedersund, Department of Biomedical Engineering
Linköping University,
SE-581 83 Linköping, Sweden.


### Supplementary Figures

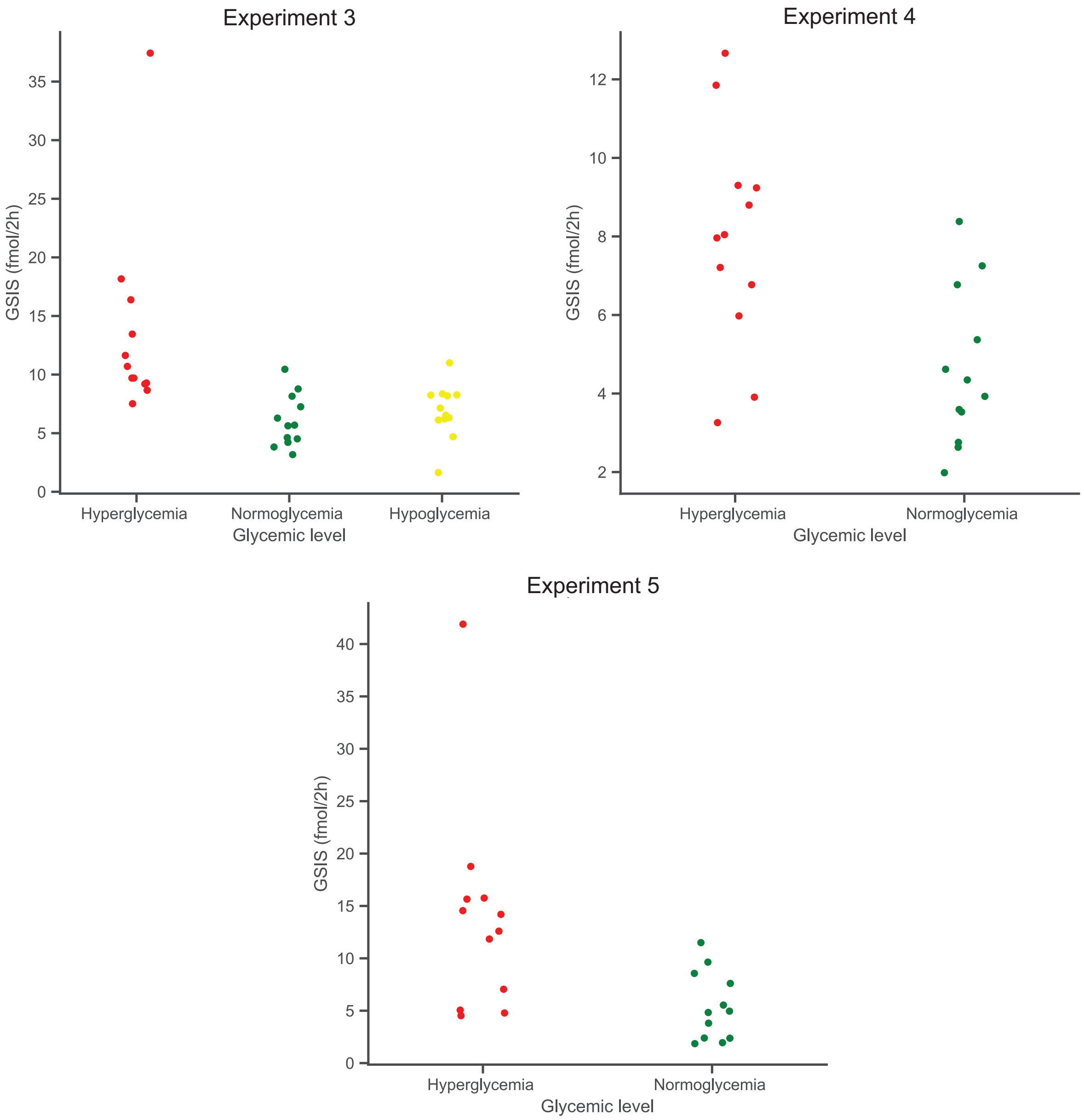


**Figure S1: Comparison between glucose-stimulated insulin secretion (GSIS) from pancreatic islets exposed to different glycemic levels during the co-culture.** The pancreatic islets were collected from the MPS after 15 days of co-culture. During the co-culture, they were exposed to either hyper- (11mM, red), normo- (5.5 mM, green) or hypoglycemic conditions (2.8 mM, yellow). At day 13, a GTT with a glucose load of 11 mM was performed in all co-cultures. After being collected from the MPS, the pancreatic islets were incubated in low glucose (2.8 mM) over 2h, following 2h incubation in high glucose (16.8 mM). The results correspond to experiments 3 (A), 4 (B) and 5 (C).

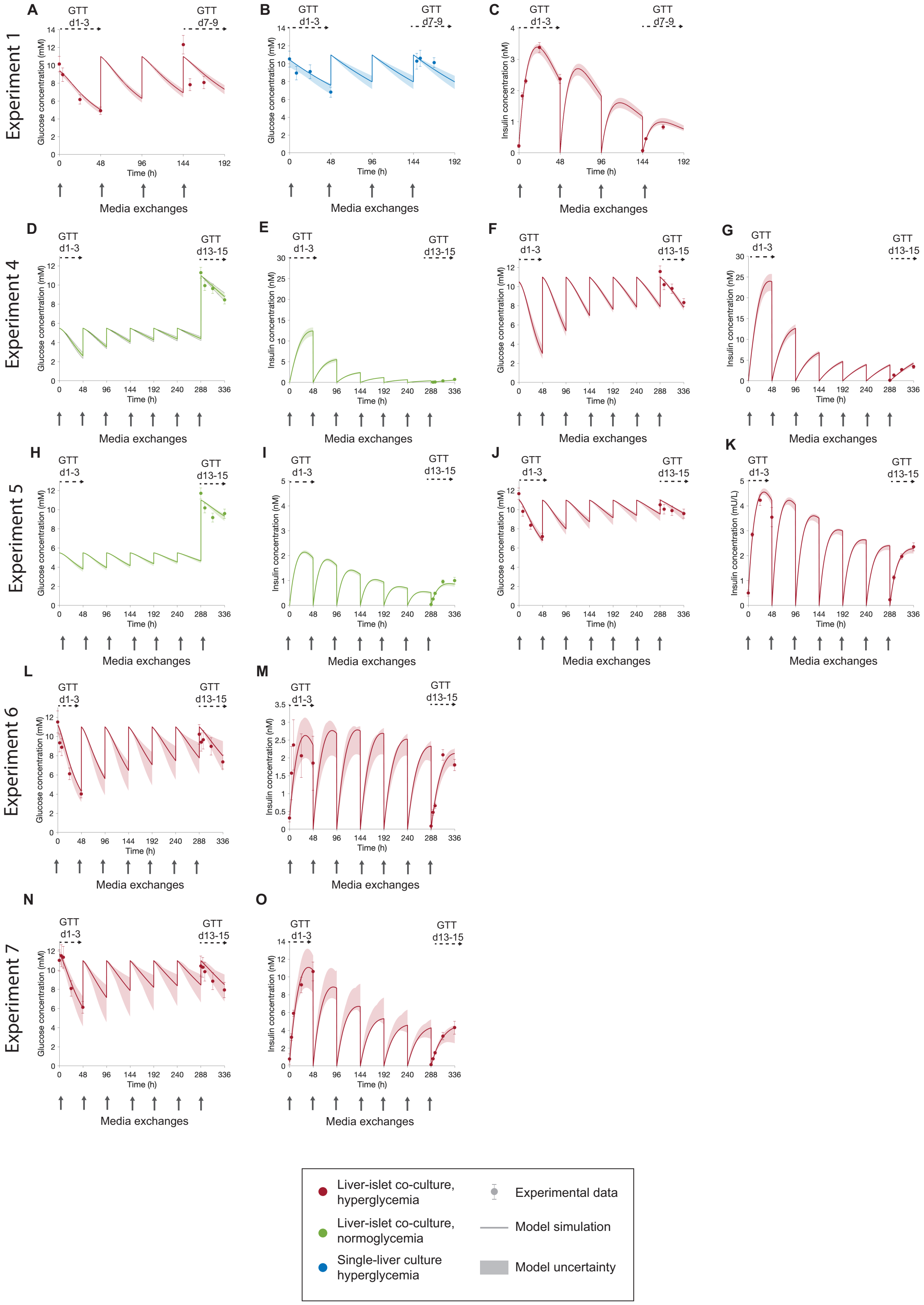


**Figure S2: Agreement between model simulations (lines) and experimental data (markers) for the experiments not shown in the main article (experiments 1 and 4-7). Experiment 1 (A-C):** Glucose concentration in the liver-islet co-culture under hyperglycemia **(A)**, glucose concentration in the single-liver culture under hyperglycemia **(B)**, insulin concentration in the liver-islet co-culture under hyperglycemia **(C)**;

**experiment 4** (**D-G**): glucose concentration in the liver-islet co-cultures under normoglycemia (**D**), insulin concentration in the liver-islet co-culture under normoglycemia (**E**), glucose concentration in the liver-islet co-culture under hyperglycemia (**F**), insulin concentration in the liver-islet co-culture under hyperglycemia (**G**); **experiment 5** (**H-K**): glucose concentration in the liver-islet co-culture under normoglycemia (**H**), insulin concentration in the liver-islet co-culture under normoglycemia (**I**), glucose concentration in the liver-islet co-culture under hyperglycemia (**J**), insulin concentration in the liver-islet co-culture under hyperglycemia (**K**); **experiment 6** (**L-M**): Glucose concentration in the liver-islet co-culture under normoglycemia (**L**), insulin concentration in the liver-islet co-culture under normoglycemia (**M**); **experiment 7** (**N-O**): Glucose concentration in the liver-islet co-culture under hyperglycemia (**N**), insulin concentration in the liver-islet co-culture under hyperglycemia (**O**). In hyperglycemic and normoglycemic conditions, co-cultures were exposed to 11 mM or 5.5 mM glucose in each media exchange (arrows), respectively. Model uncertainty is shown as shaded areas in panels A-O. Data in panels A-O are presented as mean ± SEM, where the number of replicas considered for each experiment are: n=4 (experiments 1, 6 and 7 for all glycemic conditions), n=5 (experiment 4 for all glycemic conditions and experiment 5 for hyperglycemia) and n=10 for experiment 5 under normoglycemia.

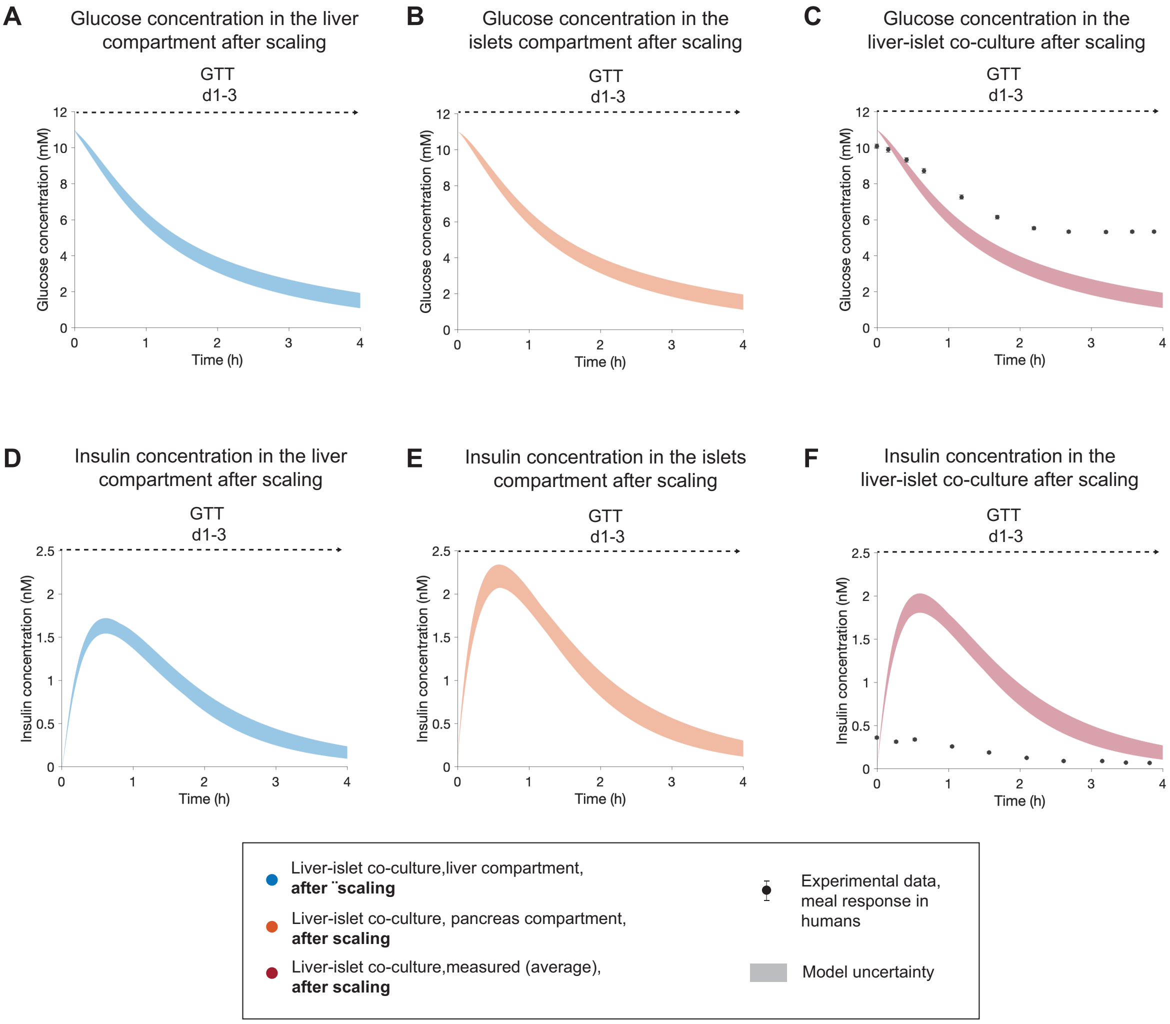


**Figure S3: Model predictions of glucose and insulin concentrations in the MPS after scaling to humans, considering the initial secretion rate of the β cells estimated in the co-culture (σmax = 6·10⁶ mIU/L/h).** The results correspond to a single experiment (experiment 1). **A,B,D,E**: Model predictions of glucose **(A,B)** and insulin **(D,E)** in the liver and pancreas compartments after scaling. **C** shows the comparison between the model prediction of plasma glucose concentration after scaling and experimental data of glucose response to a meal in healthy subjects (Man et al., 2007). The model-based prediction of the insulin response and the experimental measurements of insulin are compared in **F**. The predictions are computed for the GTT initiated at day 1 (GTT d1-3). The experimental data were acquired in a group of 204 normal subjects (Man et al., 2007). We consider the time point of peak glucose concentration in the experimental data as time = 0 h for this study, since the MPS lacks an intestinal compartment and glucose is administered directly to both the liver and pancreas compartments. Data are presented as mean ± SEM (n=204). Model uncertainty is depicted as shaded areas in (**A-F**).

### Supplementary Tables

**Table S1: Summary of the experimental settings used in the MPS experiments and the measurements acquired for calibration and evaluation of the experiment-specific computational models.** GTT: Glucose tolerance test.

| Experiment number | Experiment arms | Length of co-culture | Glycemic regimes | Time of GTTs | Measurements for the experiment-specific computational model |
|---|---|---|---|---|---|
| 1[1] | Single-liver<br>Liver-islet | 9 days | Hyperglycemia | Single-liver: Days 1-3, Days 7-9<br>Liver-islet: Days 1-3, Days 7-9 | **Single-liver:** Glucose concentration during GTTs (calibration)<br>**Liver-islet:** Glucose and insulin concentrations during GTTs (calibration) |
| 2[1] | Single-liver<br>Liver-islet | 15 days | Hyperglycemia | Single-liver: Days 1-3, Days 13-15<br>Liver-islet: Days 1-3, Days 13-15 | **Single-liver:** Glucose concentration during GTTs (calibration)<br>**Liver-islet:** Glucose and insulin concentrations during GTTs (calibration); Glucose and insulin concentrations at both liver and pancreas compartments measured every 48 h between days 3 and 13 of the co-culture (evaluation) |

[1] Experiments included in (Bauer et al., 2017)

| | | | | | |
|---|---|---|---|---|---|
| 3 | Liver-islet | 15 days | Hyperglycemia<br>Normoglycemia<br>Hypoglycemia | Hyperglycemia: Days 1-3, Days 13-15<br>Normoglycemia: Days 13-15<br>Hypoglycemia: Days 13-15 | Glucose and insulin concentrations during GTTs for hyper-and normo-glycemia (calibration)<br>Glucose and insulin concentrations during GTTs for hypoglycemia (evaluation) |
| 4 | Liver-islet | 15 days | Hyperglycemia<br>Normoglycemia | Days 13-15 | Glucose and insulin concentrations during GTTs for all glycemic regimes (calibration) |
| 5 | Liver-islet | 15 days | Hyperglycemia<br>Normoglycemia | Hyperglycemia: Days 1-3,Days 13-15<br>Normoglycemia: Days 13-15 | Glucose and insulin concentrations during GTTs for all glycemic regimes (calibration) |
| 6 | Liver-islet | 15 days | Hyperglycemia | Days 1-3, Days 13-15 | Glucose and insulin concentrations during GTTs |
| 7 | Liver-islet | 15 days | Hyperglycemia | Days 1-3, Days 13-15 | Glucose and insulin concentrations during GTTs |

**Table S2: Summary of model evaluations for all the MPS experiments included in the analysis:** Acceptable models could be inferred for every experiment. The threshold for the $\chi^2$ test with 95% significance is calculated based on the number of data points in the experimental time-series for each experiment.

| Experiment number | Agreement with experimental data ($\chi^2$ error, threshold) |
|---|---|
| 1 | 29.45 (<31.41) |
| 2 | 21.62 (<31.41) |
| 3 | 7.55 (<37.65) |
| 4 | 17.48 (<26.27) |
| 5 | 28.37 (<37.65) |
| 6 | 20.21 (<31.41) |
| 7 | 7.01 (<31.41) |

**Table S3: Estimated values of the experiment-specific parameters for each MPS experiment included in the study.** Values indicated as (-) were not estimated because there were no available data to perform the estimation.

| **Parameter** | **Unit** | **Estimated value for each MPS experiment** | | | | | | |
|---|---|---|---|---|---|---|---|---|
| | | **1** | **2** | **3** | **4** | **5** | **6** | **7** |
| $E_{G0}$ | 1/h | 1.19 | 1.03 | 1.05 | 0.75 | 0.36 | 0.83 | 0.84 |
| $CL_{I,spheroids}$ | 1/h | 17.35 | 15.56 | 22.57 | 1.24 | 11.77 | 7.04 | 6.55 |
| $S_{I0}$ | L/mIU/h | $3.12 \cdot 10^{-3}$ | $2.47 \cdot 10^{-3}$ | $1.08 \cdot 10^{-2}$ | $1.63 \cdot 10^{-3}$ | $4.33 \cdot 10^{-3}$ | $1.68 \cdot 10^{-2}$ | $2.79 \cdot 10^{-3}$ |
| $\sigma_{max}$ | mIU/L/h | $6.85 \cdot 10^{6}$ | $1.08 \cdot 10^{7}$ | $5.31 \cdot 10^{6}$ | $1.70 \cdot 10^{7}$ | $5.60 \cdot 10^{6}$ | $2.64 \cdot 10^{6}$ | $9.68 \cdot 10^{6}$ |
| $\alpha$ | $h^2$ | 87.76 | 181.09 | 194.42 | 48.07 | 157.89 | 383.92 | 107.54 |
| $k_v$ | | 1.93 | 10 | 13.60 | 26.97 | 10.20 | 1 | 16.46 |
| $I_{max,Si}$ | | $1.81 \cdot 10^{-3}$ | 0.57 | 1 | 0.53 | 0.99 | 0.98 | 0.97 |
| $EC50_{Si}$ | mmol·h/L | 4621.79 | 100.01 | 100.1 | 138.55 | 195.61 | 100 | 100.2 |
| $\Delta G_{d1}$ | mmol/L | -1.78 | -2 | - | -0.5 | -0.04 | -0.78 | 0.62 |
| $\Delta G_{d13}$ | mmol/L | - | -1.55 | - | 0.42 | -0.28 | -0.94 | -0.67 |
| $\Delta I_{d1}$ | mIU/L | 34.49 | 34.38 | - | 0.15 | 90.95 | 49.94 | 160.64 |
| $\Delta I_{d13}$ | mIU/L | - | 0 | - | 21.71 | 32.63 | 10.85 | 23.61 |

### Supplementary description of the computational model: model equations

The computational model developed in this study is formulated with the set of ordinary differential equations (ODEs) described below. A detailed description of the key equations in the model is given in the main article (Methods, Section 2.2).

#### *Glucose dynamics in the liver compartment*

Glucose in the co-culture medium of the liver compartment is determined by the dose of glucose, glucose production and uptake from the liver spheroids, and glucose exchange with the pancreas compartment:

$$\frac{dNG_{m,liver}(t)}{dt} = G_d(t) + V_{HepaRG,hep} \cdot EGP(t) - V_{hep}\left(E_{G0} + S_I(t) \cdot \frac{NI_{m,liver}(t)}{V_{m,liver}}\right)\frac{NG_{m,liver}(t)}{V_{m,liver}} + Q \cdot \frac{NG_{m,pancreas}(t)}{V_{m,pancreas}} - Q \cdot \frac{NG_{m,liver}(t)}{V_{m,liver}} \ (mmol/h)$$

$NG_{m,liver}(t)$: Number of glucose molecules in the liver compartment (mmol)

$NG_{m,pancreas}(t)$: Number of glucose molecules in the pancreas compartment (mmol)

$NI_{m,liver}(t)$: Number of insulin molecules in the liver compartment (mIU)

$G_d(t)$: Glucose input rate to the liver and pancreas compartments, determined by glucose variations because of media exchanges. Co-culture medium with 11 mM, 5.5 mM or 2.8 mM glucose is added into each of the culture compartments in each media exchange for the hyper-, normo- and hypoglycemic regimes, respectively.

$E_{G0}$: Insulin-independent glucose disposal rate of the liver spheroids (1/h)

$EGP(t)$: Endogenous glucose production from the liver spheroids (mmol/L/h)

$Q$: Flow rate between culture compartments (L/h)

$V_{HepaRG,spheroids}$: Volume of HepaRG cells in the liver spheroids (L)

$V_{m.liver}$: Volume of co-culture media in the liver compartment (L)

$V_{m.pancreas}$: Volume of co-culture media in the pancreas compartment (L)

The insulin sensitivity of the liver spheroids, $S_I(t)$, is described as follows:

$$S_I(t) = S_{I0} \cdot \left(1 - \frac{I_{max,Si} \cdot G_{int}(t)}{EC50_{Si} + G_{int}(t)}\right) \; (L/mIU/h)$$

$S_{I0}$: Insulin sensitivity of the liver spheroids at the start of the co-culture (L/mIU/h)

$I_{max,Si}$: Maximal fractional reduction of insulin sensitivity

$EC50_{Si}$: Value of time integral of excess glucose (i.e. difference between glucose levels in the co-culture media and the normoglycemic concentration of 5.5 mM) providing half of $I_{max,Si}$ (mmol·h/L)

The variable $G_{int}(t)$ increases progressively as the liver spheroids are exposed to glucose levels above the normoglycemic range $\left(\frac{NG_{m,liver}(t)}{V_{m,liver}} - Gnormo \geq 0\right)$, as given by:

$$\frac{dG_{int}(t)}{dt} = \begin{cases} \frac{NG_{m,liver}(t)}{V_{m,liver}} - G_{normo} & \frac{NG_{m,liver}(t)}{V_{m,liver}} - Gnormo \geq 0 \\ 0 & \frac{NG_{m,liver}(t)}{V_{m,liver}} - Gnormo < 0 \end{cases} \quad \left(\frac{mmol}{L}\right)$$

$G_{normo}$: Glucose concentration considered for normoglycemia (5.5 mM)

#### *Glucose dynamics in the pancreas compartment*

Glucose content in the pancreas compartment is described as:

$$\frac{dNG_{m,pancreas}(t)}{dt} = G_d(t) + Q \cdot \frac{NG_{m,liver}(t)}{V_{m,liver}} - Q \cdot \frac{NG_{m,pancreas}(t)}{V_{m,pancreas}} \; (mmol/h)$$

#### *Insulin dynamics in the liver compartment*

Insulin content in the liver compartment is described by the following equation:

$$\frac{dNI_{m,liver}(t)}{dt} = Q \cdot \frac{NI_{m,pancreas}(t)}{V_{m,pancreas}} - Q \cdot \frac{NI_{m,liver}(t)}{V_{m,liver}} - V_{HepaRG,spheroids} \cdot CL_{I,spheroids} \cdot \frac{NI_{m,liver}(t)}{V_{m,liver}} \; (mIU/h)$$

$CL_{I,spheroids}$: Insulin clearance by the liver spheroids (1/h)

*Insulin dynamics in the pancreas compartment*

Insulin content in the pancreas compartment is determined by the release of insulin from the $\beta$ cells in the pancreatic islets:

$$\frac{dNI_{m,pancreas}(t)}{dt} = V_{\beta,islets}(t) \cdot \sigma(t) \cdot \frac{\left(\frac{NG_{m,pancreas}(t)}{V_{m,pancreas}}\right)^2}{EC50_I{}^2 + \left(\frac{NG_{m,pancreas}(t)}{V_{m,pancreas}}\right)^2} + Q\frac{NI_{m,liver}(t)}{V_{m,liver}} - Q\frac{NI_{m,pancreas}(t)}{V_{m,pancreas}} \ (mIU/h)$$

$EC50_I$ : Glucose concentration resulting in half-of-maximum response to insulin (mM)

The insulin secretion capacity per unit volume of $\beta$ cell is given by:

$$\sigma(t) = \sigma_{max} \cdot \left(1 - \frac{t^2}{\alpha \quad + t^2}\right) (mIU/L/h)$$

$\sigma_{max}$: Maximal insulin secretion capacity per unit volume of $\beta$ cell (i.e. at the start of the co-culture) (mIU/L/h)

$\alpha$ : Parameter defining the sigmoidal dependence on time of $\sigma(t)$ ($h^2$)

The variable $V_{\beta,islets}(t)$ (L) describes the changes in volume of $\beta$-cells in the pancreatic islets over the co-culture time, according to the following equation:

$$\frac{dV_{\beta,islets}(t)}{dt} = k_v(-d_0 + r_1 G_{slow,pancreas}(t) - r_2 G_{slow,pancreas}(t)^2) \cdot V_{\beta,islets}(t) \ \ (L/h)$$

where $d_0$ is the death rate at zero glucose ($h^{-1}$) and $r_1 = r_{1,r} + r_{1,a}$ (L/mmol/h) and $r_2 = r_{2,r} + r_{2,a}$ ($L^2/mmol^2/h$), where $r_{1,r}$, $r_{1,a}$ (L/mmol/h), $r_{2,r}$, $r_{2,a}$ ($L^2/mmol^2/h$) are parameters that determine the dependence of the replication and apoptosis rates on glucose. The parameter $k_v$ was introduced to account for potential differences in behaviour between pancreatic islets in our *in vitro* system and rodent islets the model of Topp et al. (Topp et al., 2000).

The variable $G_{slow,pancreas}(t)$ (mmol/L) represents the long-term average (i.e daily) glucose concentration in the co-culture mediumas given by:

$$\frac{dG_{slow,pancreas}(t)}{dt} = \frac{G_{pancreas}(t) - G_{slow,pancreas}(t)}{\tau_{slow}} \ (mmol/L/h)$$

$G_{pancreas}(t)$ : Glucose concentration in the pancreas compartment (mmol/L)

$\tau_{slow}$ : Time constant that determines the averaging of $G_{pancreas}(t)$ over time (h)

The concentrations of glucose and insulin in each compartment were calculated by dividing the insulin and glucose content, respectively, by the volume of co-culture medium in the compartment:

$$G_{liver}(t) = \frac{NG_{m,liver}(t)}{V_{m,liver}} \text{ (mmol/L)}$$

$$G_{pancreas}(t) = \frac{NG_{m,pancreas}(t)}{V_{m,pancreas}} \text{ (mmol/L)}$$

$$I_{liver}(t) = \frac{NI_{m,liver}(t)}{V_{m,liver}} \text{ (mIU/L)}$$

$$I_{pancreas}(t) = \frac{NI_{m,pancreas}(t)}{V_{m,pancreas}} \text{ (mIU/L)}$$

Glucose and insulin samples in the MPS were obtained by pooling samples from both the liver and the pancreas compartment. Therefore, the resulting glucose and insulin measurements ($G(t)$ and $I(t)$, respectively), were computed as:

$$G(t) = {G_{liver}(t) \cdot V_{sample,liver} + G_{pancreas}(t) \cdot V_{sample,pancreas}}\Big/{(V_{sample,liver} + V_{sample,pancreas})} \text{ (mmol/L)}$$

$$I(t) = {I_{liver}(t) \cdot V_{sample,liver} + I_{pancreas}(t) \cdot V_{sample,pancreas}}\Big/{(V_{sample,liver} + V_{sample,pancreas})} \text{ (mIU/L)}$$

where $V_{sample,liver}$ and $V_{sample,pancreas}$ are the volumes of co-culture media collected from the liver and pancreas compartment in each sample (15 µl).

The initial conditions for the model states are listed below:

$NG_{m,liver}(0) = (G_{dose} + \Delta G_{d1}) \cdot V_{m,liver}$ (mmol)

$NG_{m,pancreas}(0) = (G_{dose} + \Delta G_{d1}) \cdot V_{m,islets}$(mmol)

$NI_{m,liver}(0) = \Delta I_{d1} \cdot V_{m,liver}$ (mIU)

$NI_{m,pancreas}(0) = \Delta I_{d1} \cdot V_{m,pancreas}$ (mIU)

$t(0) = 0$ (h)

$G_{int}(0) = 0$ (mmol·h/L)

$G_{slow,pancreas}(0) = 5.5$ (mmol/L)

$V_{\beta,islets}(0) = 8.8 \cdot 10^{-9}$ (L)

where $\Delta G_{d1}$ (mmol/L), $\Delta I_{d1}$ $(mIU/L)$ are offset parameters that account for experimental errors related to the media exchange performed at day 1. This can be due to, for instance, variations in volume of the co-culture medium or added glucose and insulin concentrations when the media exchange is performed. Similarly, the parameters $(\Delta G_{d13}, \Delta I_{d13})$ were included in the model to account for these errors during the media exchanges performed at day 13.
